# Tumor -Associated MUC1 Regulates TGF-β Signaling and Function in Pancreatic Ductal Adenocarcinoma

**DOI:** 10.1101/2020.04.29.068577

**Authors:** Priyanka Grover, Sritama Nath, Mukulika Bose, Alexa J. Sanders, Cory Brouwer, Nitika, Ru Zhou, Mahboubeh Yazdanifar, Mohammad Ahmad, Shu-ta Wu, Andrew W. Truman, Pinku Mukherjee

## Abstract

Pancreatic ductal adenocarcinoma (PDA) is one of the most lethal human cancers. Transforming Growth Factor Beta (TGF-β) is a cytokine that switches from a tumor-suppressor to a tumor promoter throughout tumor development, by a yet unknown mechanism. Tumor associated MUC1 (tMUC1) is aberrantly glycosylated and overexpressed in >80% of PDAs and is associated with poor prognosis. The cytoplasmic tail of MUC1 (MUC1-CT) interacts with other oncogenic proteins promoting tumor progression and metastasis. We hypothesize that tMUC1 levels regulate TGF-β functions in PDA *in vitro* and *in vivo*. We report that high-tMUC1 expression positively correlates to TGF-βRII and negatively to TGF-βRI receptors. In response to TGF-β1, high tMUC1 expressing PDA cells undergo c-Src phosphorylation, and activation of the Erk/MAPK pathway; while low tMUC1 expressing cells activate the Smad2/3 pathway, enhancing cell death. Correspondingly, mice bearing tMUC1-high tumors responded to TGF-β1 neutralizing antibody *in vivo* showing significantly retarded tumor growth. Analysis of clinical data from TCGA revealed significant alterations in gene-gene correlations in the TGF-β pathway in tMUC1 high versus tMUC1 low samples. This study deepens our understanding of tMUC1-regulated TGF-β’s paradoxical function in PDA and establishes tMUC1 as a potential biomarker to predict response to TGF-β-targeted therapies.

## Introduction

Pancreatic Cancer is currently the third leading cause of cancer-related deaths in the United States (http://pancreatic.org/). It has been projected to become the second leading cause of cancer-related deaths in the US, surpassing colorectal cancer by the year 2020 (http://pancreatic.org/). About 93% of pancreatic cancers are Pancreatic Ductal Adenocarcinomas (PDA) with patients demonstrating a median survival rate of less than six months and a five-year survival rate of 9% in the US [1]. Worldwide, the overall five-year survival rate ranges from 2% to 9% [2]. It has a mortality rate (2.61%) that nearly matches its incidence rate (3.2%) [3].

The transforming growth factor beta (TGF-β) signaling pathway belongs to a large superfamily that primarily consists of TGF-β (including cytokines such as TGF-β1, 2, and 3), bone morphogenetic proteins, activins, and inhibins [4]. This family of growth factors activates many biological signals, such as cell growth, apoptosis, differentiation, immune response, angiogenesis, and inflammation [5–7]. Deregulation of the TGF-β pathway can lead to cancer, among other ailments [8]. In normal environments and early cancers, TGF-β1 regulates epithelial cells as a tumor suppressor by controlling the cell cycle and inducing apoptosis. However, once the cancer is established, a switch occurs and TGF-β1 becomes a tumor promoter. TGF-β1 induces invasion and migration and eventually leads to epithelial-to-mesenchymal transition (EMT) [9]. This process helps facilitate the migration and invasion of cancer cells to distant locations leading to metastasis, the major cause of cancer-related deaths [10].

Canonical TGF-β signaling is initiated by the binding of a TGF-β cytokine to a pair of specific transmembrane receptors, TGF-βRI and TGF-βRII [11]. This activates the cytoplasmic serine/threonine kinase domains of the TGF-β receptors [12], which leads to further activation downstream. In normal environments, TGF-β1 binds to its specific receptors TGF-βRII and TGF-βRI, in sequence. This leads to the phosphorylation of Smad2/3 via the cytoplasmic Serine/Threonine kinase domain of TGF-βRI [13]. Smad2 has been identified as a tumor suppressor and mediator of the antiproliferative TGF-β1 and activin responses [14]. Smad 2/3 trilocalizes with Smad4. This leads the heterotrimer complex to the nucleus to induce transcriptional changes that influence cell regulation.

Frequent alterations and changes in the TGF-β pathway occur in cancer. Misregulated TGF-β signaling activates Erk1/2 leading to an increase in aggressive cancer characteristics, such as growth, invasion, migration, and metastasis [15]. Smad4 alterations are also particularly common in PDA. Smad4 is inactivated in about half of PDA cases, where homozygous deletions account for 30% of PDA cases and loss of heterozygosity accounts for 20% of all PDA cases [16].

Mucin-1 (MUC1) is a Type I transmembrane glycoprotein that influences tumor progression and metastasis in PDA [17]. Tumor-associated MUC1 (tMUC1) is overexpressed and aberrantly glycosylated in more than 80% of PDA cases [17–21]. In normal environments, MUC1 is expressed on the apical surface of ductal cells to provide a protective barrier [22]. However, upon tumorigenesis MUC1 expression is no longer restricted to the apical surface. At this point, MUC1 glycosylation decreases and the protein becomes overexpressed across the cell surface, placing it into the close vicinity of many growth factor receptors [19]. tMUC1 oncogenic signaling, which plays an important role in increased metastasis and invasion, is promoted through the cytoplasmic tail (tMUC1-CT). The tMUC1-CT is a highly conserved 72-amino acid long section containing seven tyrosine residues that are phosphorylated by tyrosine receptor kinases, such as c-Src [23, 24]. Interestingly, C-Src has been shown to mediate breast cancer cell proliferation and invasion via regulation of mitogen-activated protein kinase (MAPK) activation [25]. In our previous studies, we have demonstrated that tMUC1 modulates TGF-β signaling in cell lines that were engineered to express high tMUC1 [26]. In that report, we established that TGF-β signaling required tyrosine phosphorylation of the tMUC1-CT via tyrosine kinase, c-Src. Here we deepen our understanding of tMUC1 regulation of TGF-β signaling in PDA cells that are genetically varied and that express varying levels of endogenous tMUC1. We establish that tMUC1 plays a definitive role in switching TGF-β1 from a tumor suppressor to a tumor promoter. In the presence of high tMUC1, TGF-β activates the Erk pathway, promotes c-Src phosphorylation, and enhances resistance to apoptosis. In PDA cells with low levels of tMUC1, TGF-β activates the Smad pathway, decreases c-Src phosphorylation, and induces cell death. Taken together, our work suggests a novel reciprocal interaction between tMUC1 and TGF-β. The *in vivo* data suggest that tMUC1 may be a potential biomarker for future anti-TGF-β targeted therapies in PDA.

## Results

### High expression of tMUC1 in PDA cells positively correlates to TGF-βRII and negatively correlates to TGF-βRI levels, leading to activation of Erk pathway

Several studies have established that overexpression of tMUC1 in PDA is linked to enhanced growth and metastasis [21, 27, 28]. To investigate a possible correlation between tMUC1 and TGF-β signaling we selected a panel of human and mouse PDA cell lines with varying levels of tMUC1, Smad4, and p53 expression (Fig. 1A) and assessed the expression of tMUC1, TGF-β RI, TGF-β RII by Western Blotting (Fig. 1B). Cells expressing high levels of tMUC1 displayed high levels of TGF-β RII and low levels of TGF-βRI, whereas cells expressing low levels of tMUC1 had high levels of TGF-β RI and low levels of TGF-β RII. There was a significant positive correlation (0.7 with a p value of <0.01) between tMUC1 and TGF-β RII levels and a significant negative correlation (0.5 with a p value of <0.05) between tMUC1 and TGF-β RI levels (Fig. 1C).

**Figure 1.**
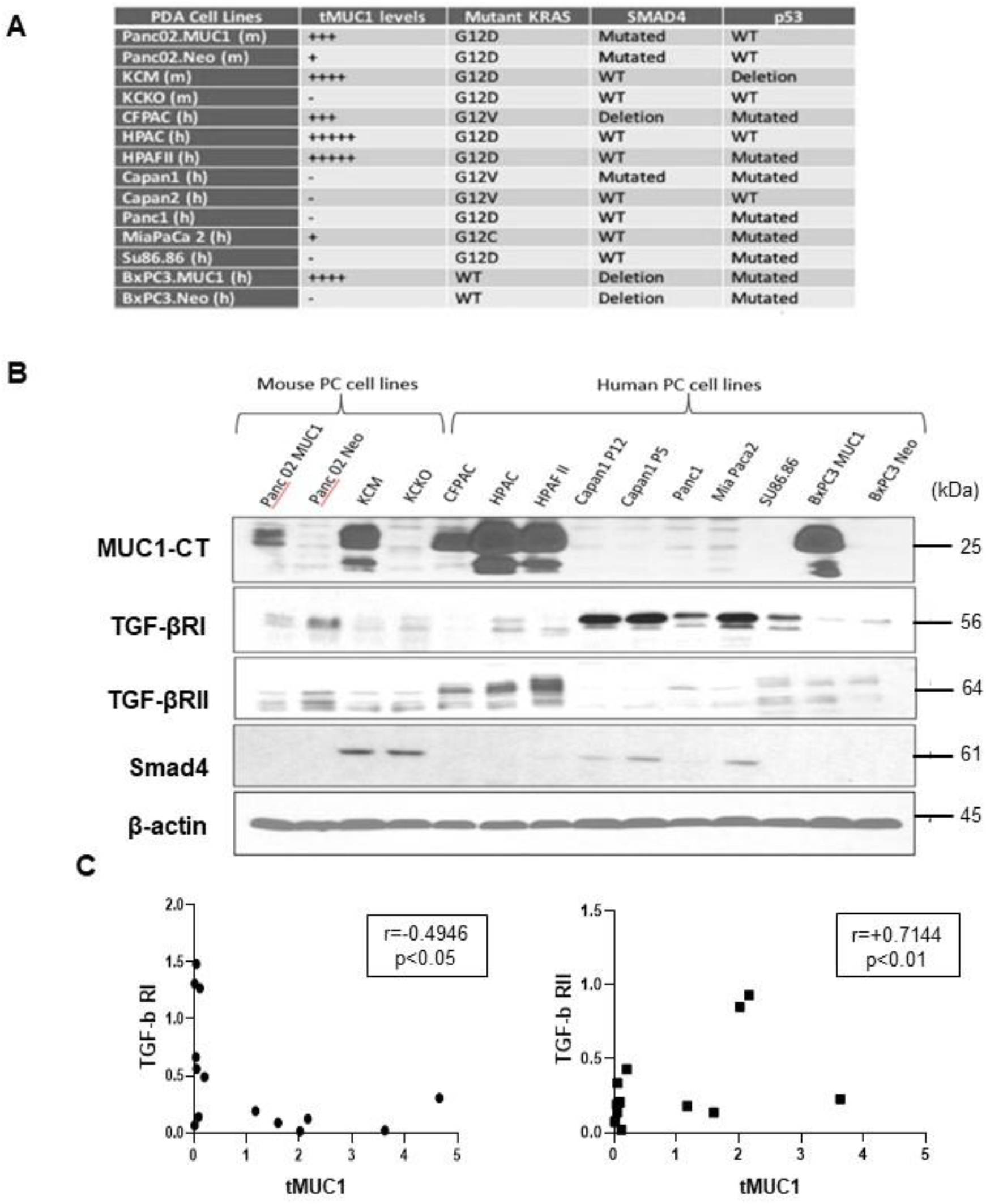
tMUC1 high expression positively correlates to TGF-βRII and negatively correlates to TGF-βRI levels. **A.** List of PDA cell lines used in the experiments with mutation status of KRAS, SMAD4 and p53 along with their tMUC1 expression levels. **B.** Western blot detecting expression of tMUC1-CT, TGF-βRI, TGF-βRII, and SMAD4 in a panel of PDA cell lines. β-actin was used as an endogenous loading control. **C.** Densitometric analysis of tMUC1 expression compared to TGF-βRI expression shows a significantly negative correlation between the two (Pearson’s correlation coefficient r=-0.4946, p<0.05). **D.** Densitometric analysis of tMUC1 expression compared to TGF-βRII expression shows a significantly positive correlation between the two (Pearson’s correlation coefficient r=0.7144, p<0.01).

To determine if signaling downstream is affected by the differences in TGF-β RI and II levels, we examined changes in phosphorylation of Erk1/2 and Smad2/3 over time to 10ng/ml of TGF-β (Fig. 2A). For this experiment, we used three tMUC1 high (HPAF-II, HPAC, and CFPAC) and three tMUC1 low (MiaPaCa-2, Panc1, and BxPC3 WT) cell lines. We observed that two out of three cell lines with high tMUC1 (HPAF II and CFPAC) showed increased phosphorylation of Erk1/2 (Fig. 2A) in response to TGFβ1 in 5, 15, and 30 minutes. Interestingly, the low tMUC1 cells (MiaPaCa2, Panc1 and BxPc3) showed high p-Erk1/2 at 0 time and decreased in 5- and 10-minutes post TGF-β1 stimulation but this recovered by 30 minutes of exposure (Fig. 2A). In contrast, almost all cell lines started out with low to undetectable levels of pSmad2/3. Post TGF-β1 treatment, we observed a steady increase in p-Smad2/3 in two of the three tMUC1 low cells (Panc1 and BxPC3) while the two tMUC1 high cells (HPAFII and CFPAC) showed undetectable levels of pSmad2/3 even at 30 minutes (Fig. 2A).

**Figure 2.**
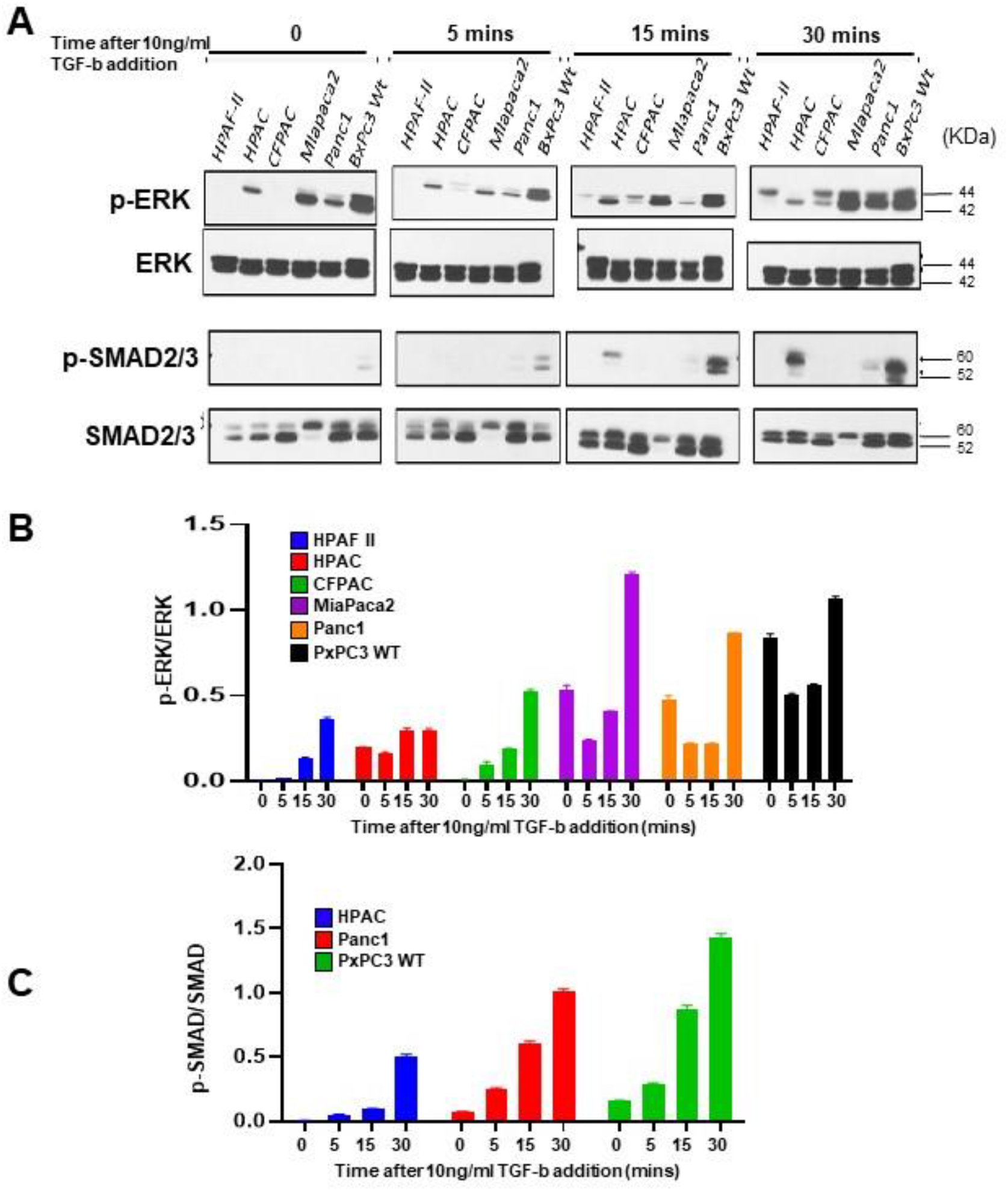
High tMUC1 PDA cells respond to TGF-β1 by phosphorylation of Erk1/2 and low tMUC1 by phosphorylation of SMAD2/3. **A.** Western blot expression of phosphorylation of Erk1/2 and Smad2/3 compared to total Erk1/2 and Smad2/3 in a panel of PDA cells in response to TGF-β1 at 0, 5, 15, and 30 minutes. **B.** Corresponding densitometric analysis of p-Erk1/2 in all PDA cell lines is presented. Arbitrary densitometric unit of p-Erk1/2 normalized to total Erk1/2. **C)** Corresponding densitometric analysis of p-SMAD2/3 in all PDA cell lines expressing p-SMAD2/3 is presented. Arbitrary densitometric unit of p-SMAD2/3 normalized to total-SMAD2/3. Ratios of p-Erk1/2 normalized to total-Erk1/2 and p-SMAD2/3 normalized to total-SMAD2/3 are presented as means +/− SEM of n=3. * p < 0.05, ** p < 0.01, *** p<0.001, **** p<0.0001.

Fascinatingly, in spite of high tMUC1 in HPAC cells, the p-Erk1/2 remained unchanged while the pSmad2/3 increased in response to TGFβ1 by 15 and 30 minutes (Fig. 2A, 2B, and 2C). This contrast to other tMUC1 high PDA cell lines may be possibly explained by the presence of a wild type p53 protein in HPAC cells. There is an established crosstalk between TGF-β and p53 signaling [29]. The mutational status of p53 has been reported to interfere with TGF-β functioning, thus switching it from tumor suppressive to protumorigenic [30]. MiaPaca2 is the only cell line to have a G12C mutation in KRAS and it has been shown that mutant KRAS-G12D has higher affinity for PI3K, whereas mutant KRAS-G12C has higher affinity for Ras-Ral guanine nucleotide dissociation stimulator (RalGDS) [31]. This might result in a constitutive phosphorylation and activation of Erk in MiaPaca2 cells. MiaPaca2 cells are TGF-β R II negative [32] and are non-responsive to TGF-β exposure, so no phosphorylation of SMAD2/3 is expected. Some low-tMUC1 PDA cell lines express steady-state levels of p-SMAD2 in absence of TGF-β [33]. The data suggests that the high tMUC1 cells use the noncanonical (Erk1/2) pathway in response to TGF-β1 stimulation (possibly via engaging TGF-β RII), whereas the low tMUC1 cells use the canonical (Smad2/3) pathway via engaging TGF-β RI.

### TGF-β1 exposure increases cell death in low tMUC1 cells

We next investigated the effects of TGF-β1 treatment on the cell cycle in high and low tMUC1 cells. To this end, we exposed high (HPAFII and CFPAC) and low (MiaPaCa2 and BxPC3) tMUC1 cells to TGF-β1 for 48 hours. There were minimal differences in the cell cycle phases (G2M, S, G1/G0, and sub G0) in the tMUC1 high cells post treatment with TGF-β1 (Fig. 3A) suggesting that high tMUC1 cells may be less sensitive on TGF-β1 for mitosis or cell death. In contrast, we observed a significant increase in Sub-G0 phase in both the MiaPaca2 and Panc1 cells in response to TGF-β1 treatment (Fig. 3B). Interestingly, we also observed a significant decrease in G1 and G2M phase in MiaPaca2 and Panc1 cells respectively (Fig. 3B). Taken together, the data conforms our hypothesis that tMUC1 low cells may respond to TGF-β1 by slowing down mitosis and increasing cell death.

**Figure 3.**
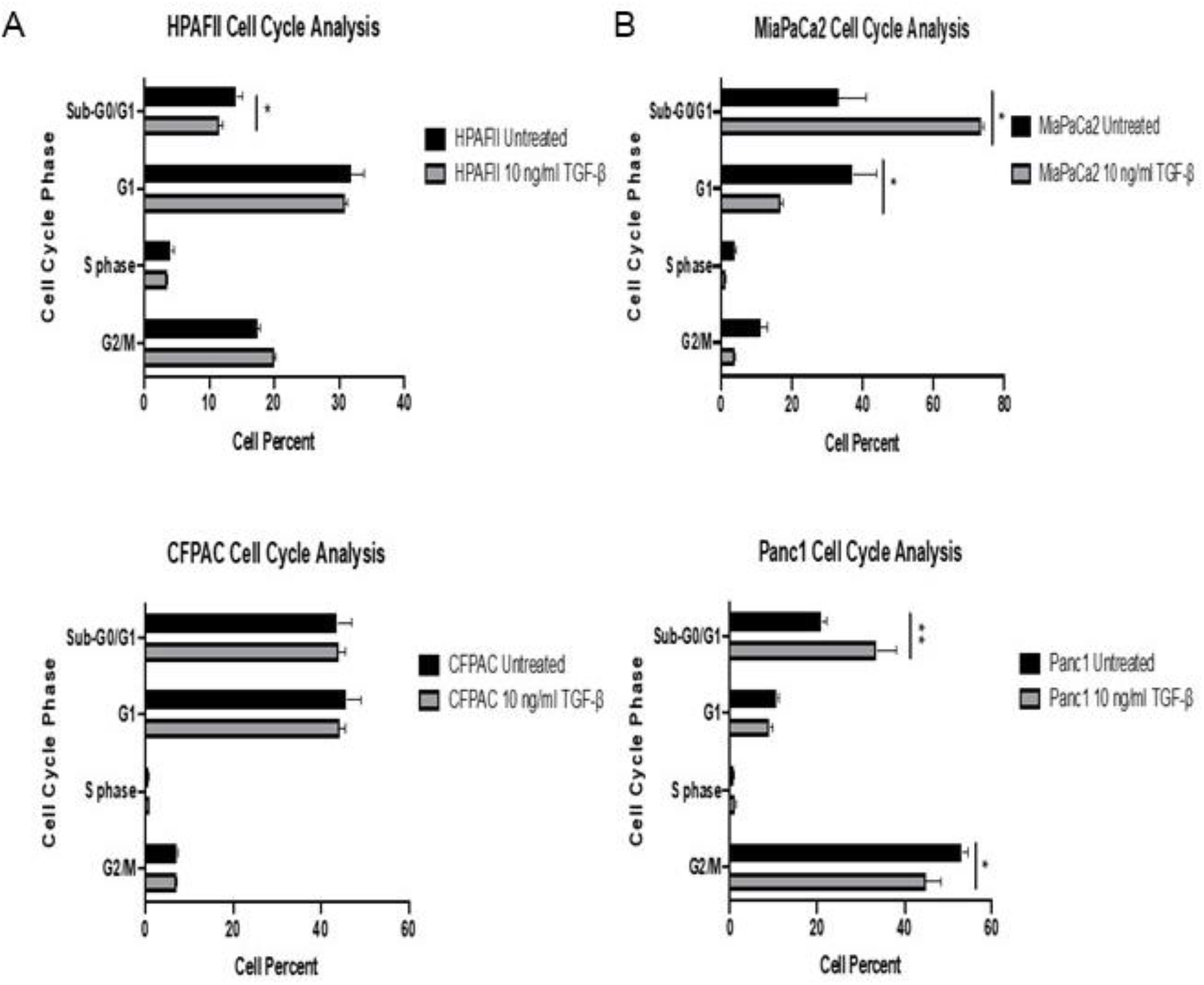
TGF-β1 exposure increases cell death in low tMUC1 PDA cells. Cell cycle analyses of **A.** tMUC1 high (HPAF II and CFPAC) and **B.** tMUC1 low (MiaPaca2 and Panc1) cells in response to 10 ngs/ml of TGF-β1. All data are shown as bar graphs as means +/− SEM of n=3. * p < 0.05, ** p < 0.01, *** p<0.001, **** p<0.0001.

### Increased phosphorylation of c-Src in tMUC1-high cells in the response to TGF-β1 treatment

Our previous studies established c-Src as the key mediator in the interaction of tMUC1 and TGF-β signaling [34]. Therefore, we examined the phosphorylation of c-Src in HPAF-II, CFPAC, and MiaPaca2 cells in response to 10 and 50ng/ml of TGF-β1 treatment for 30 minutes. SRC kinases are regulated by phosphorylation events-for example, c-Src has an autophosphorylation site at Y416 that induces activation and a regulatory site at Y527 which is phosphorylated by Csk and Chk that induce c-Src inhibition [35] [36]. Phosphorylated Y527 stabilizes a closed conformation of c-Src, which suppresses kinase activity, while phosphorylated Y416 promotes kinase activity of c-Src by stabilizing its activation loop to facilitate substrate binding [37]. After 30 minutes of TGF-β1 exposure, we observed increased phosphorylation of c-Src at Y416 (Fig. 4A) and decreased phosphorylation at Y527 (Fig. 4B), both of which signify activation of c-Src kinase activity. Interestingly, in the low tMUC1, MiaPaCa-2 cells, we observed just the opposite, decreased phosphorylation at Y416 and increased phosphorylation at Y527 suggesting dephosphorylation of c-Src in MiaPaca2 cells in response to TGF-β1 treatment.

**Figure 4.**
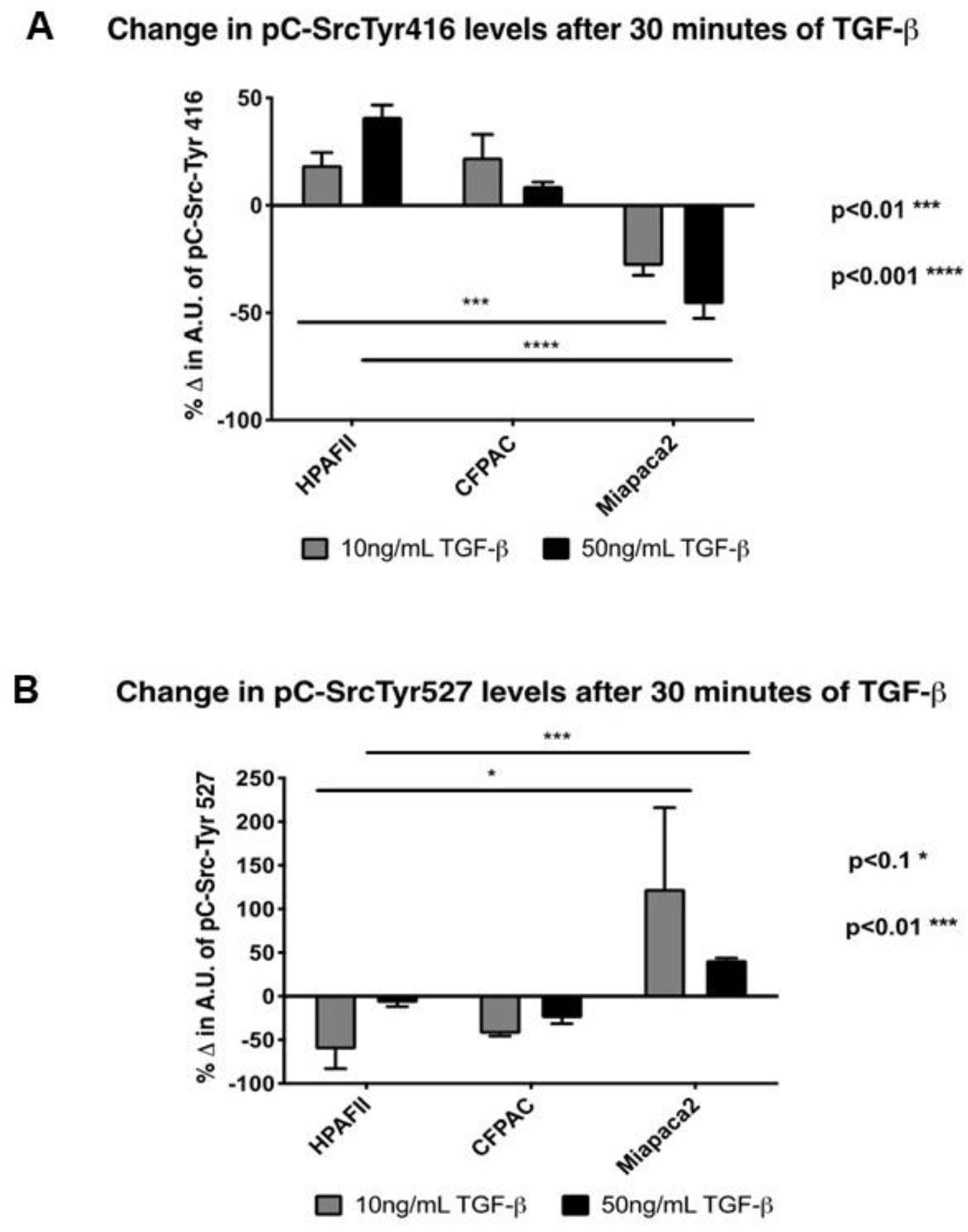
c-Src activation in high tMUC1 cells in response to TGF-β1. After 30 minutes of exposure to 10 and 50 ngs/ml of TGF-β1, c-Src phosphorylation at **A.** Tyr416 and **B.** Tyr527 was tested by western blotting and densitometric analysis conducted to generate the OD values. Percent change from untreated O.D. values are illustrated. First, each density unit for the particular protein was normalized to their respective β-actin density. Percent change was determined by formula ((TGF-β treated – No treatment)/No treatment) x 100. If the final answer was negative, this was percentage decrease (suggesting that the protein level remained unchanged with treatment). All densitometry OD values were normalized to total c-Src protein. Densitometric analysis is shown as means +/− SEM of n=3. * p < 0.05, ** p < 0.01, *** p<0.001, **** p<0.0001. One-way ANOVA comparing the data across the different cell lines.

### TGF-β regulates tMUC1 protein expression via proteasomal degradation

While our data suggested tMUC1 modulation of TGF-β signaling, we also considered the intriguing possibility that the reverse may be true-that TGF-β signaling regulates tMUC1 expression. Imbalance of protein homeostasis is known to cause cellular malfunction and lead to diseases including tumor growth and metastasis [38]. Therefore, we determined the effect of TGF-β1 treatment on the signaling moiety (the CT) of tMUC1. In this experiment, we elected to use engineered BxPC3 PDA cells to keep all genetic factors (other than tMUC1) the same. We compared BxPC3.MUC1 (engineered to express high levels of full-length MUC1) to BxPC3.Neo (engineered to express the empty vector), and BxPC3.Y0 (engineered to express full-length MUC1 with all 7 tyrosines of the MUC1-CT mutated to phenylalanine). We treated these cells with 10ng/ml of TGF-β1 to determine the effects on tMUC1 protein expression and degradation (Fig. 5). Upon 30 minutes of treatment with TGF-β1, there was a sharp decrease in tMUC1 levels, however, when proteasomal degradation was blocked with MG132 inhibitor, the basal level of tMUC1 was fully restored in BxPC3.MUC1 cells (Fig. 5A and 5B). This was not as evident in the low tMUC1 BxPC3.Neo cells (Fig. 5A and 5B). Interestingly, we observed no change in protein expression in the BxPc3.Y0 cells suggesting that the degradation pathway depends on tyrosine phosphorylation of tMUC1-CT (Fig. 5A and 5B). Lomako *et al* (2009) established that TGF-β regulates MUC4 in MUC4-transfected melanoma cells by repressing precursor cleavage. Proteasome inhibitors were shown to repress the TGF-β inhibition of MUC4 expression [39]. As before, TGF-β1 treatment resulted in a decrease in tMUC1-CT protein expression. However, using MG132 (a proteasome inhibitor), we were able to rescue ~90% of the degradation, suggesting that TGF-β1 regulates tMUC1 levels through proteasomal degradation. Furthermore, degradation was dependent upon the presence of the 7 tyrosine residues of MUC1-CT, overall establishing TGF-β signaling as a novel regulator of tMUC1 in PDA.

**Figure 5.**
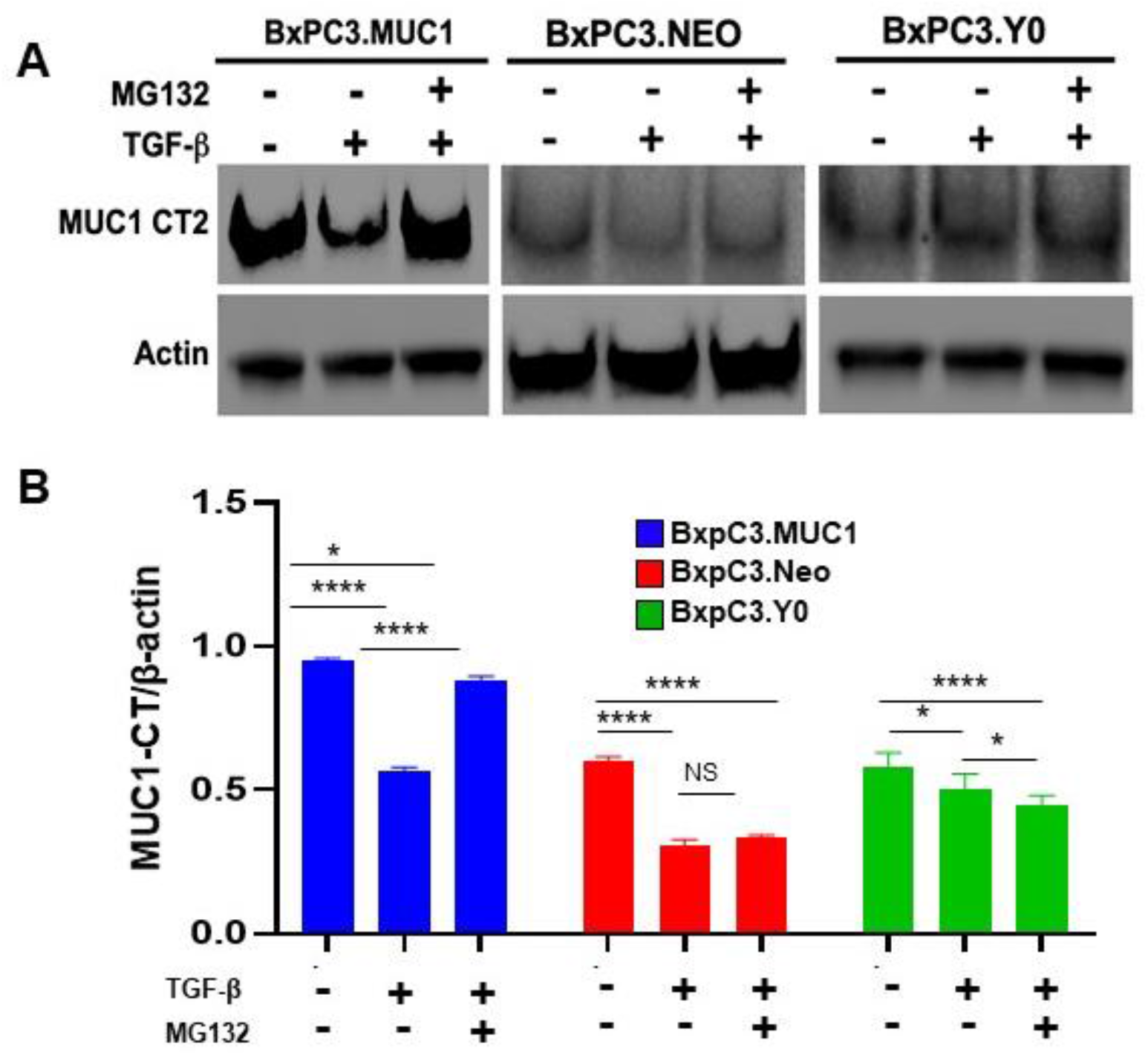
TGF-β1 promotes the degradation of tMUC1. **A.** BxPC3.MUC1, BxPC3.Neo and BxPC3.Y0 cells were treated with either 10 ngs/ml of TGF-β alone or in combination with 100 μM MG132 for 30 minutes. Immunoblotting was used to evaluate the tMUC1 levels using the CT2 antibody. β-actin was used as loading control. **B.** Densitometric analysis of tMUC1 normalized to β-actin are presented as means +/− SEM of n=3. * p < 0.05, ** p < 0.01, *** p<0.001, **** p<0.0001.

### Neutralizing TGF-β antibody treatment significantly dampens tMUC1 high tumor growth but has no effect on tMUC1 low tumors in vivo

Given that our data shows high expression of tMUC1 in PDA promotes hyperactivation of Erk pathway signaling in response to TGF-β1, we hypothesized that treatment with anti-TGF-β1 neutralizing antibody would hamper growth of tMUC1 high but not low tMUC1 tumors *in vivo*. Athymic Nude-Foxn1nu mice bearing tMUC1-high (HPAF-II) or tMUC1-low (MiaPaCa2) established tumors were injected intra-tumorally with either control IgG or neutralizing TGF-β antibody three times a week for two weeks (Fig. 6A). The established tumors were measured with calipers over 28 days and once euthanized, tumor wet weight was taken. tMUC1 high (HPAF II) tumors had significantly lower tumor growth and tumor wet weight when treated with TGF-β1 antibody as compared to the control IgG group (Fig. 6A and Fig. 6C). In contrast, tMUC1 low (MiaPaca2) tumors did not respond to TGF-β1 neutralizing antibody treatment (Fig. 6B and D). In fact, TGF-β1 antibody treated MiaPaca2 tumors weighed slightly more (albeit not significant) than the IgG treated group (Fig. 6D). The treatment did not have any adverse effect on the weight or well-being of the mice (Supplemental Fig. S1).

**Figure 6.**
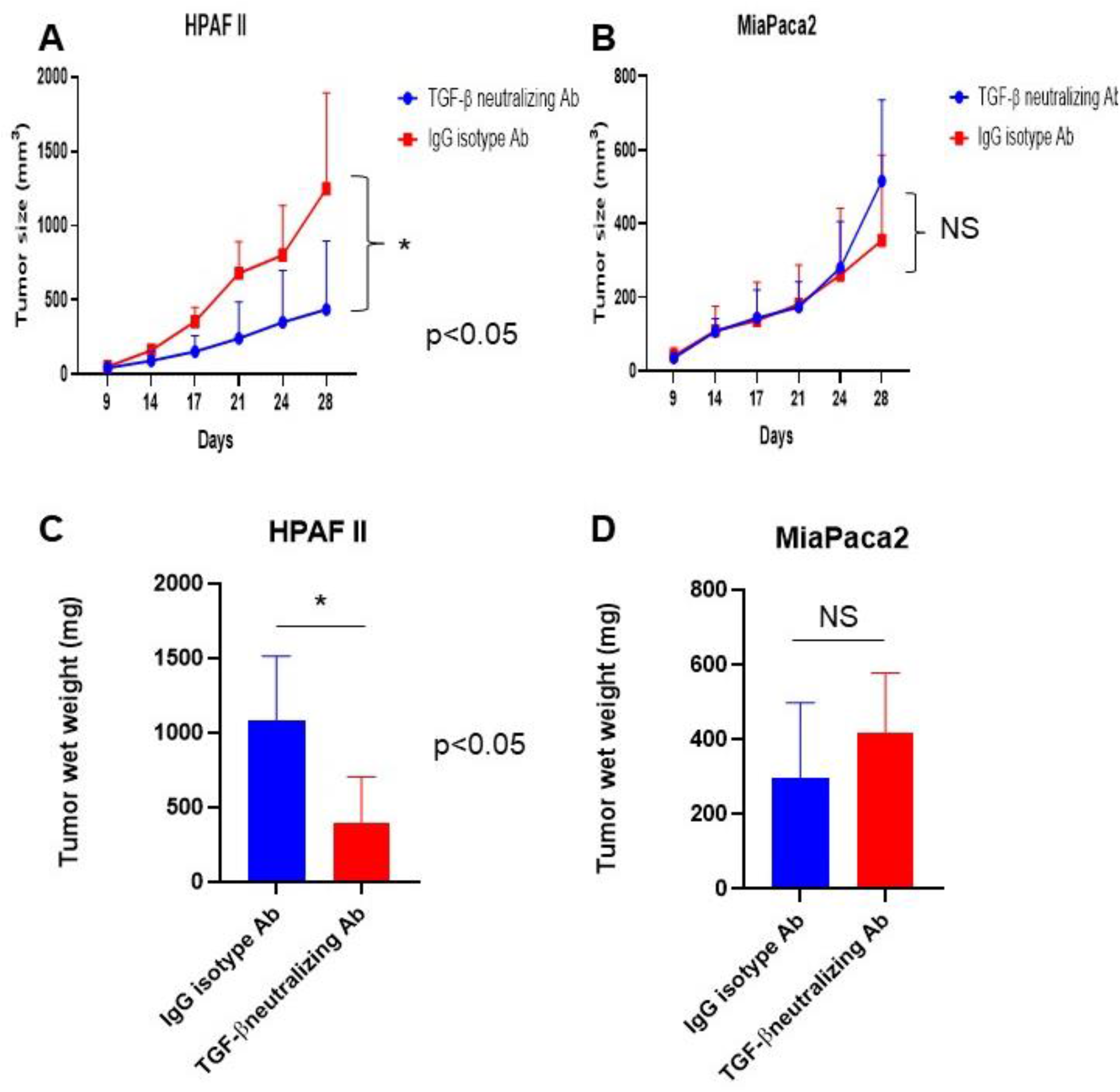
Neutralizing TGF-β1 antibody treatment significantly reduced tMUC1 high (HPAF II) but not tMUC1 low (MiaPaca2) tumor growth *in vivo*. Tumor growth of **A.** HPAFII (n=5 for TGF-β neutralizing Ab and n=4 for IgG isotype)and **B.** MiaPaca2 (n=6 for both groups) measured using caliper measurements weekly and tumor size in mm^3^ is plotted. **C. and D.** Tumor wet weight of HPAFII and MiaPaca2 tumors respectively. *p<0.05, NS: non-significant.

### Distinct gene correlations based on high versus low tMUC1 in PDA patient samples

To assess the clinical significance of our results, we utilized the available TCGA database and generated a heatmap to demonstrate the correlation between each of the genes in two groups: low tMUC1 expression samples and moderate/high tMUC1 expression samples (Fig. 7). Genes were prefiltered based on a defined list containing genes involved in the TGF-β pathway. Both groups were compared to normal samples to establish if the gene correlation was stronger or weaker in either low tMUC1 or moderate/high tMUC1 samples. We found that SMAD4 had strong negative correlation to high versus low tMUC1 samples. More importantly, correlation of SMAD4 to CDKN2B, ID4, SRC, RHOA, RAF1 was distinctly different in high versus low tMUC1 samples. Similarly, correlation of ID4 to RHOA, RAF1 and SRC are also distinct between tMUC1 high versus low samples. These data corroborate the clinical significance of high versus low tMUC1 PDAs and that TGF-β signaling predominantly promote oncogenic signal in high tMUC1 PDA while favoring an apoptotic signaling in the low tMUC1 PDA. The correlation of these six genes to expression of MUC1 alone is depicted in Fig. 7B. The changes in correlation pattern of these genes with MUC1 in high versus low-MUC1 expressing samples are further illustrated in Fig. 7C.

**Figure 7.**
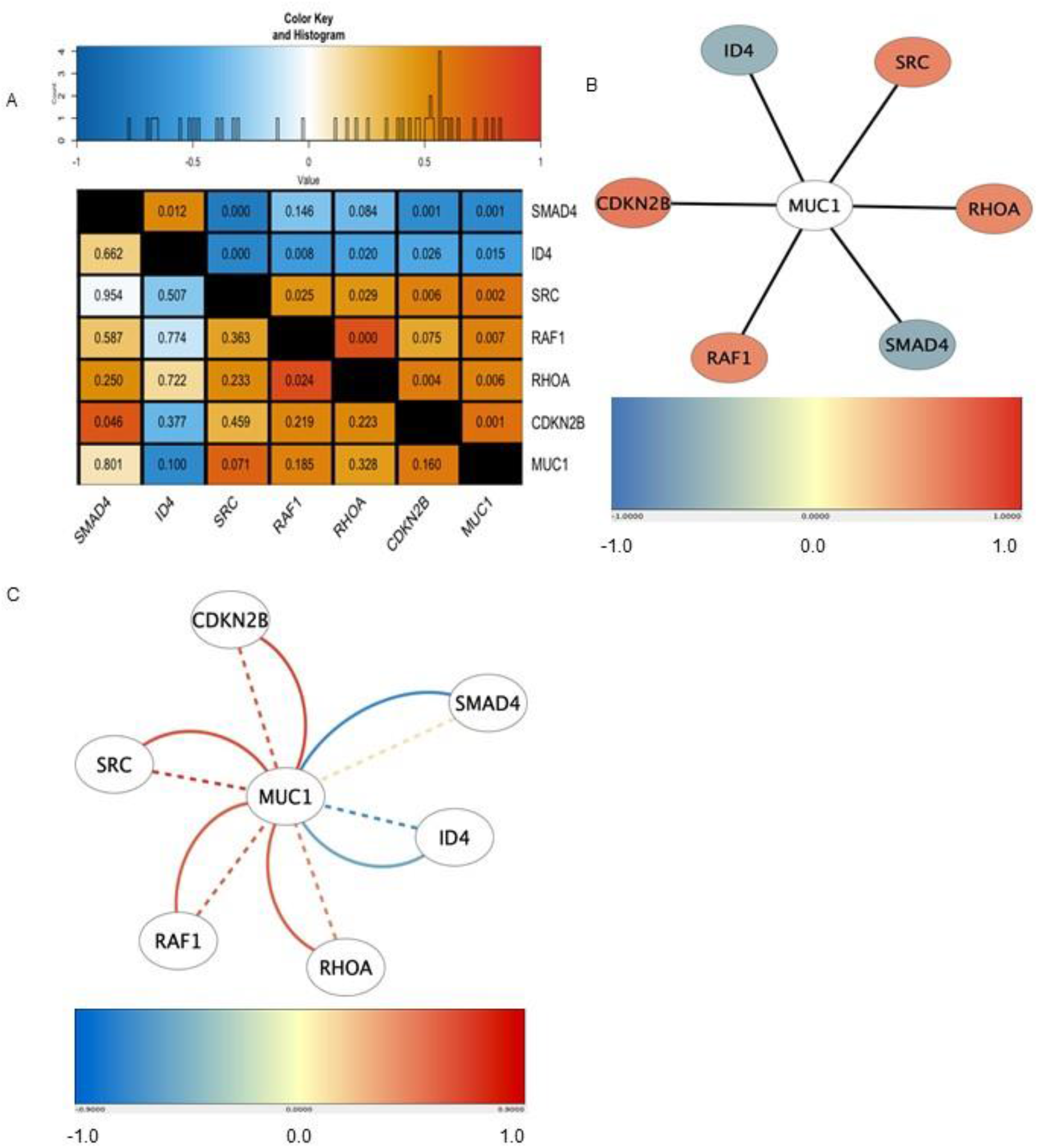
TCGA correlation heatmap showing important gene correlations in high and low MUC1 PDA samples. **A.** Top panel shows the color key histogram of correlation values. Bottom panel shows the heatmap generated showing correlation between tMUC1 to significant genes of the TGF-β pathway in 29 PDA tumor samples. The tMUC1 horizontal and vertical rows show the correlation of each gene with tMUC1 in the low and high tMUC1 expression samples respectively. Comparing the color block of each gene with itself in the tMUC1 row versus column is a visual demonstration of the correlation change between the genes dependent on tMUC1 expression. The lower left triangle shows gene correlations in low tMUC1 expression samples and the upper right triangle shows gene correlations in the moderate/high tMUC1 expression samples. Respective p-values for each gene-gene correlation are mentioned in the boxes. **B.** This figure shows the genes that are significantly correlated with tMUC1 in the 29 tumor samples. The color of the nodes (circles with gene names) are the correlation values of each gene with tMUC1 on a scale from –1 to 1. **C.** This figure shows genes significantly correlated with tMUC1 in tumor samples, separated by tMUC1 expression. The color of the dashed lines corresponds to the correlation value between each gene with tMUC1 in the low tMUC1 expression group. The color of the solid lines corresponds to the correlation value between each gene with tMUC1 in the moderate/high tMUC1 expression group. Genes with a false discovery rate adjusted p<0.05 are shown.

## Discussion

In 2009, the National Cancer Institute of NIH chose tMUC1 as the second most targetable antigen in adenocarcinomas [40]. tMUC1 is overexpressed in more than 80% of PDA cases [17] and TGF-β signaling plays an important oncogenic role in majority of cancers including in PDA [9]. The data presented demonstrates that tMUC1 regulates TGF-β signaling and function via activation of c-Src and in turn TGF-β1 modulates tMUC1 proteasomal degradation via signaling through the 7 tyrosines of the tMUC1-CT. The data has important clinical relevance because tMUC1 expression can be used as a surrogate biomarker to determine the efficacy of future TGF-β-targeted treatments.

Using a panel of PDA cell lines, we demonstrate that high tMUC1 expression is positively correlated to TGF-βRII expression (Fig. 1), phosphorylation of Erk1/2 (Fig. 2), and activation of c-Src (Fig. 4); all of which signify activation of the non-canonical pathway associated with cellular proliferation and invasion [34] [41]. Furthermore, high tMUC1 negatively correlated to TGF-βRI expression, a receptor that activates the canonical Smad pathway, (Fig. 1) known to drive cells towards cell death and apoptosis [42]. Our hypothesis that tMUC1 regulates TGF-β signaling and function was further confirmed in low-tMUC1 expressing PDA cells. As was expected, low tMUC1 expression positively correlated to TGF-βRI expression, activation of the canonical Smad 2/3 pathway (Fig. 2), and cell death (Fig. 3).

Thus far, we were focused on the role of tMUC1 in regulation of TGF-β signaling and function. In this study we show that TGF-β1 may play a significant role in MUC1 proteasomal degradation (Fig. 5). To the best of our knowledge, this is the first evidence of TGF-β1 involvement in MUC1 degradation and its dependence on the 7 tyrosines in MUC1-CT. In future experiments, we will delineate which of the 7 tyrosines (or combination) may be critical for the TGF-β1 induced degradation of MUC1. In our previous publication, we found no difference in tMUC1 expression levels after 48 hours of TGF-β1 exposure [34] which may have been too late to detect differences. Therefore, we determined that TGF-β1 affects tMUC1 levels immediately (30 minutes) post treatment. Although regulation of proteasomal degradation of MUC4 by TGF-β1 was established in melanoma cells by repressing precursor cleavage [39], this is the first such report on MUC1 degradation in PDA. It has been shown that protein degradation by proteasomes leads to small peptides that can yield biologically active protein fragments, such as transcription factors NF-kB and Mga2p [43]. Very little has been uncovered regarding tMUC1 degradation and this novel finding will be further studied for future publication. The *in vitro* data signifies an intricate crosstalk between tMUC1 and TGF-β signaling and oncogenesis. It is plausible that a constant barrage of TGF-β1 leads to tMUC1 degradation in cancer cells, ultimately creating active protein fragments that induces oncogenic gene expression.

If indeed TGF-β signaling is critical in the aggressive growth of tMUC1-high PDA tumors, then neutralizing TGF-β1 with an antibody *in vivo* in mice would dampen tumor growth. Indeed, neutralizing TGF-β1 treatment in tMUC1-high HPAFII tumors significantly reduced tumor progression and reduced tumor burden (Fig. 6A and C), whereas, the same treatment had no effect on tMUC1-low MiaPaca2 tumors (Fig. 6B and D).

Lastly, we analyzed PDA cases from TCGA database based on tMUC1 expression levels (Fig. 7). The correlation heatmap corroborated some of *in vitro* data but also uncovered novel correlations in tMUC1 high versus low PDAs. Correlation of SMAD4 with all other genes (ID4, SRC, RAF1, RHOA, CDKN2B and MUC1) was clearly distinct in high versus low tMUC1 PDA (Fig. 7A) with a strong negative correlation with tMUC1 in high-tMUC1 samples (Fig. 7C). SMAD4 is a member of the SMAD family of signal transducers and is a central mediator of TGF-β signaling pathways [44]. The SMAD-dependent TGF-β pathway is the canonical pathway and is known to promote tumor suppression by inducing cell cycle arrest, apoptosis, and maintenance of genomic integrity. However, in most PDA, TGF-β loses its tumor suppressive function often by the inactivation of SMAD4 [45]. And we see here that high expression of tMUC1 correlates significantly with SMAD4 expression in a negative manner, demonstrating the role of tMUC1 in this inhibition of tumor suppressive function of TGF-β. Inhibitor of DNA binding 4 (ID4) shows a stronger negative correlation to tMUC1 and a weaker positive correlation to SMAD4 in the low versus high tMUC1 PDA. Activation of SMAD family of genes induces expression of ID proteins, which inhibit differentiation and induce cell growth by inhibiting the function of basic helix-loop-helix transcription factors [46]. Most of the reported function for ID4 is in gastric and colorectal cancers [47] [48]. To the best of our knowledge, this is the first report of ID4’s possible role in PDA and its negative correlation to tMUC1 and positive correlation to SMAD4. In gastric cancer cell lines, ID4 is hypermethylated and infrequently expressed in gastric adenocarcinomas with high expression in normal gastric mucosa. ID4 promoter hypermethylation (leading to down regulation) significantly correlates to microsatellite instability [47]. Epigenetic inactivation of ID4 in colorectal carcinomas correlates with poor prognosis [48]. These data suggest that downregulation of ID4 leads to progression of tumor and it is known that high tMUC1 expression is a marker of poor prognosis. This makes perfect sense since ID4 and tMUC1 show negative correlation with one another in PDA.

SRC expression and activity is correlated with poor prognosis in a variety of human cancers [49] and so is the expression of tMUC1. Our correlation analysis from the TCGA data shows a positive correlation between SRC to tMUC1 in both high and low tMUC1 PDA, confirming the importance of SRC both in the canonical and non-canonical TGF-β signaling pathways. We also know that c-Src-mediates phosphorylation of the tyrosines in MUC1-CT which increases binding of MUC1 and β-catenin [50]. Nuclear co-localization of MUC1 CT with β-catenin stabilizes β-catenin in the nucleus [51] leading to transcription of EMT-associated genes in various adenocarcinomas including breast, renal, gastric and pancreas [52]. Although intriguing, we do not fully understand the high negative correlation of SRC to SMAD4 in the tMUC1-high PDA. We will be following up on this interesting finding.

RAF1, RHOA and CDKN2B all show positive correlation with tMUC1 in both high-tMUC1 and low-tMUC1 tumors (Fig. 7). However, all of these genes had a negative correlation to SMAD4 in tMUC1 high PDA while showing positive correlation to SMAD4 in tMUC1 low PDA. Data once again indicates the distinct TGF-β signaling pattern associated with tMUC1 levels in PDA, possibly leading to different outcomes. RAF1 is a proto-oncogene downstream of the RAS. KRAS mutation is known as one of the driver mutations of PDA and therefore increased expression of RAF1 in these samples and its correlation with tMUC1 denotes their oncogenic role. RHOA is a GTPase known to be highly expressed in gastric cancers [53] [54]. RHOA can be activated by TGF-β via Smad-dependent or Smad-independent pathways to induce EMT [44]. CDKN2B is an inhibitor of CDK4/6 (cyclin dependent kinase 4 and 6), which are necessary for cell cycle progression [55]. Over expression of CDKN2B inhibits progression of cell cycle leading to apoptosis [56]. Activation of canonical SMAD-dependent TGF-β signaling leads to increase in CDKN2B. Therefore, positive correlation of CDKN2B to SMAD4 in tMUC1-low PDA samples indicate cell cycle arrest and apoptosis as seen in the *in vitro* functional assay (Fig. 3) [57].

The data presented here establishes tMUC1 as a modulator of TGF-β signaling and function in PDA and determines a novel role of TGF-β1 in possibly inducing proteasomal degradation of tMUC1. A model (Fig. 8) illustrates our current understanding from this data of tMUC1’s role in switching TGF-β signaling from a tumor suppressor to a promoter.

**Figure 8.**
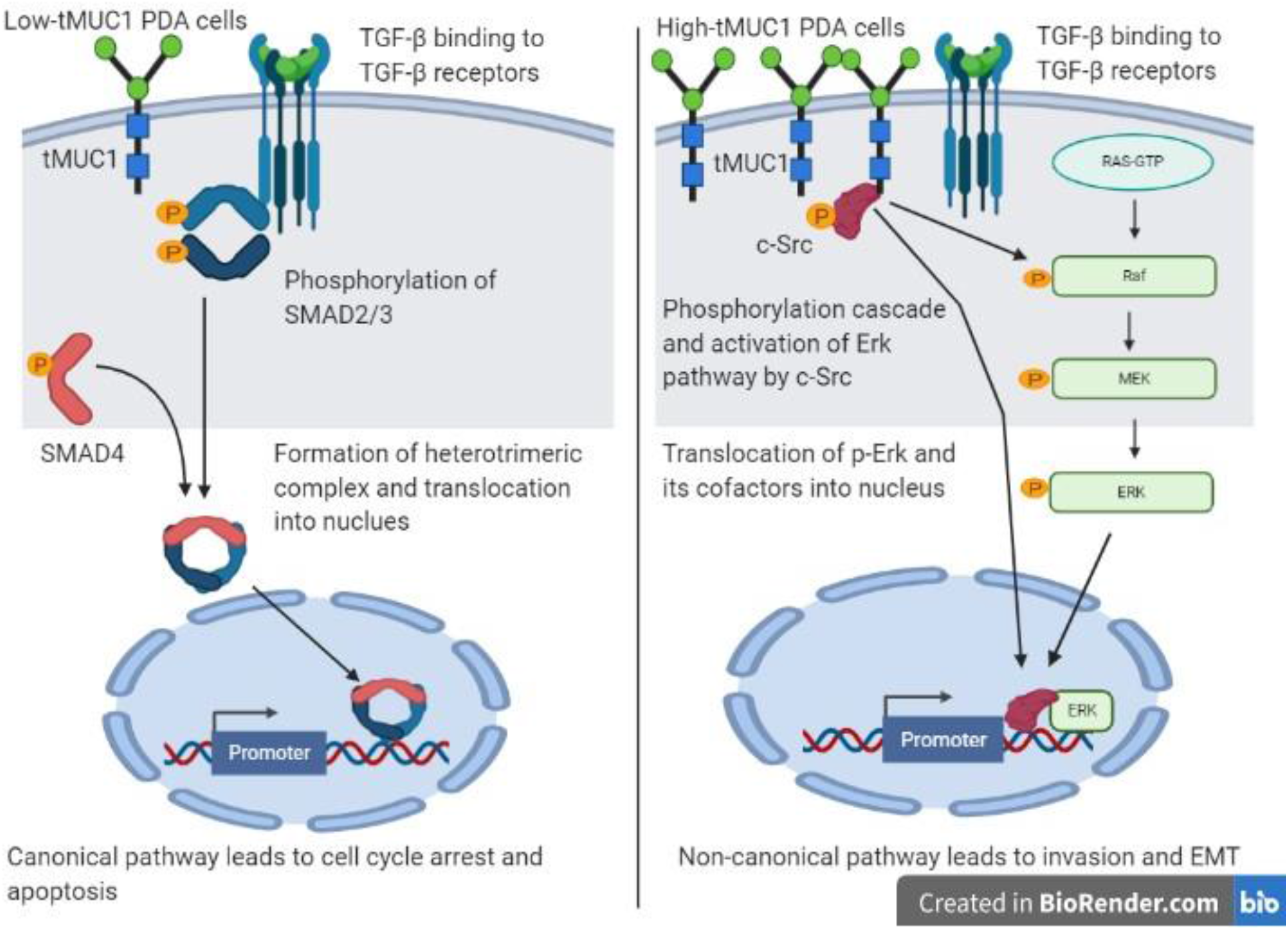
Schematic diagram of the proposed mechanism of TGF-β signaling and functions in tMUC1 high versus low PDA. **Left panel** shows activation of SMAD-dependent canonical pathway in low-tMUC1 PDA cells. TGF-β ligands bind to the membranous TGF-β receptor (TGF-βRII) homodimers with high affinity. TGF-βRII binding allows dimerization with TGF-β type I receptor (TGF-βRI) homodimers, activation of the TGF-βRI kinase domain and signal transduction via phosphorylation of the C-terminus of receptor-regulated SMADs (R-SMAD), SMAD2 and SMAD3. The TGF-βR dimer then forms a heterotrimeric complex with SMAD4 which translocates in the nucleus [62, 63]. This leads to cell cycle arrest and apoptosis of PDA cells, thus TGF-β acts as a tumor suppressor. **Right panel** shows activation of SMAD-independent non-canonical pathway in high-tMUC1 PDA cells. In this pathway, binding of TGF-β to its receptors increases phosphorylation of c-Src which in turn phosphorylates Ras and increases its activity [64]. This activates the Erk pathway and phosphorylated Erk translocates into the nucleus to increase transcription of oncogenic proteins and leads to invasion and EMT of PDA cells [65]. Thus, in high-tMUC1 PDA cells TGF-β acts as a pro-tumorigenic cytokine. The schematic was created with the basic version of www.biorender.com.

## Materials and Methods

### Cell Lines and Culture

BxPC3.MUC1, BxPC3.Neo, and BxPC3.Y0 were generated as previously described [21]. Human cell lines (CFPAC, HPAC, HPAF-II, Capan1, Panc1, MiaPaCa2, Su86.86) were obtained from American Type Culture Collection and cultured as instructed. Panc02.MUC1 and Panc02.Neo were originally gifted by Dr. Hollingsworth (University of Nebraska), and maintained in medium containing Geneticin (G418; Invitrogen, Carlsbad, CA, USA). KCM and KCKO were developed in our lab [28]. Cell lines were maintained in Dulbecco’s Modified Eagle Medium (DMEM; Gibco), Minimal Essential Media (MEM; Gibco), or Roswell Park Memorial Institute 1640 medium (RPMI; with, L-glutamine; ThermoFisher). All media was supplemented with 10% fetal bovine serum (FBS; Gibco or Hyclone), 3.4 mM L-glutamine, 90 units (U) per ml penicillin, 90 ug/ml streptomycin, and 1% Non-essential amino acids (Cellgro). RPMI was also supplemented with Geneticin. Cells were kept in a 5% CO2 atmosphere at 37 degrees Celsius. Every passage of BxPC3 transfected cells were maintained in a final concentration of 150 micrograms/ml of the antibiotic G418 (50 mg/ml) to ensure positive selection. For all experiments, cell lines were passaged no more than 10 times.

### Western Blotting

Cellular lysate preparation and Western blotting were performed as previously described [21]. The cells were divided into different treatment groups: no treatment, 10 ng/ml of TGF-β1 (Peprotech, Rocky Hill, NJ, USA), or drugs at various concentrations for 48 hours due to more pronounced signaling. 1:500 Armenian hamster monoclonal anti-human tMUC1 cytoplasmic tail (CT2) antibody was used to probe for tMUC1 in phosphate-buffered-saline-Tween 20 (PBS-T) with 5% BSA. MUC1 CT antibody CT2 was originally generated at Mayo Clinic and purchased from Neomarkers, Inc. (Portsmouth, NH) [58]. CT2 antibody recognizes the last 17 amino acids (SSLSYNTPAVAATSANL) of the cytoplasmic tail (CT) of human MUC1. Membranes were also probed with the following antibodies from Cell Signaling Technology (1:1000): Smad4 (Rabbit, 38454), p-Smad 2/3 (Rabbit, 5678), total Smad (Rabbit, 3102), p-Erk1/2 (Rabbit, 9101), total Erk (Rabbit, 9102), p-Src Family Tyr416 (Rabbit, 6943), Non-p-Src Tyr416 (Mouse, 2102), p-Src Tyr527 (Rabbit, 2105), Non-p-Src (Rabbit, 2107), and β-Actin (Mouse, 3700). Other antibodies used include TGF-βRI (Abcam, 1:200, Rabbit, ab31013) and TGF-βRII (Abcam, 1:1000, Rabbit, ab61213). Densitometric analysis was conducted using the ImageJ software and percent change was calculated accordingly: First, each density unit for the particular protein was normalized to their respective β-actin density. Percent change was determined by formula (TGF-β treated – No treatment/No treatment)x 100. If the final answer was negative, this was percentage decrease (suggesting that the protein level remained unchanged with treatment).

### For proteasomal degradation

100uM of MG132 was used to inhibit proteasomal degradation. The cells were treated for 30 minutes concurrently with TGF-β.

### Cell Cycle Analysis

To determine the effect of TGF-β1 on the cell cycle, cell lines were cultured as described above until ~80% confluent and then serum starved for 18 hours. Cells were then given 50 ng/ml of TGF-β1 in serum-free media for 24 and 48 hours. Cells were trypsinized and fixed in cold 70% ethanol in phosphate buffered saline (1x PBS) overnight. Once the cells were centrifuged, they were then stained with 50 microgram/ml Propidium Iodide (PI) with 20 microgram/ml of RNase A in 1x PBS at room temperature for 30 minutes. Approximately 30,000 cells/sample were acquired using BD LSR Fortessa flow cytometer (BD Biosciences). The data was analyzed with FLOWJO software (version 10).

### Analysis of clinical data from TCGA

Twenty-nine pancreatic adenocarcinoma RNA-Seq sample data were downloaded from the Genomic Data Commons data portal [59, 60]. All tumor samples were from the PAAD project data generated by The Cancer Genome Atlas (TCGA) Research Network: http://cancergenome.nih.gov/. The FPKM (Fragments Per Kilobase of transcript per Million mapped reads) gene expression files from these samples were analyzed to identify gene correlations with MUC1. Genes were prefiltered based on a defined list containing genes of interest, including genes involved in the TGFB pathway. Differential Gene Correlation Analysis (DGCA – version 1.0.1) package and psych_1.8.12 package in R were used to identify genes correlated with MUC1 in the tumor samples. Genes with a false discovery rate adjusted p-value of less than 0.05 were considered significantly correlated with tMUC1. The tumor samples were then separated into two groups based on their tMUC1 expression: tMUC1 low expression group and tMUC1 moderate/high expression group. A heatmap was generated for expression of significant genes and tMUC1 in the 29 tumor samples. There were 7 samples that showed clear visual low expression in the heatmap compared to the other samples; these were selected for the low expression group. The other 22 samples were included in the mod/high expression group. A heatmap showing the correlation of tumor-significant genes with tMUC1 was plotted using DGCA, separated by the tMUC1 expression groups.

### Subcutaneous Mouse Model

Athymic Nude-Foxn1nu mice were purchased from Harlan Laboratories and housed at UNC Charlotte’s vivarium. These mice were injected subcutaneously with tumor cells. 3×10^6^ HPAFII cells (50ul) (n=9) or 5×10^6^ MiaPaCa2 cells (50ul) (n=12) were injected with Matrigel (50ul) (total=100ul) subcutaneously into the flank of male or female Athymic Nude-Foxn1nu mice [61]. Once the tumors reached a palpable size (~3×3mm, ~5 days post tumor inoculation), mice were separated into 4 different groups. Groups 1 and 2 had HPAF-II tumors and groups 3 and 4 had MiaPaca2 tumors. Groups 1 and 3 were treated with the isotype control IgG antibody (20ug/100ul per mouse) three times a week for two weeks. Groups 2 and 4 were treated with the neutralizing TGF-β antibody (20ug/100ul per mouse) three times a week for two weeks. Mice were monitored daily for general health and tumors were palpated. Caliper measurements were taken three times a week until endpoint (tumor size: ~15×15mm). This study and all procedures were performed after approval from the Institutional Animal Care and Use Committee of UNC Charlotte.

### Statistical Analysis

The data are expressed as the mean +/− SEM of n=3. Differences between groups were examined using unpaired one-tailed t-tests, one-way and two-way ANOVA. Statistical comparisons were made using the GraphPad Prism 8.0. p-values of <0.05 were considered statistically significant (*p < 0.05; **p < 0.01; ***p<0.001; ****p<0.0001).

## Conclusion

The data has uncovered a major role of tMUC1 in regulating the paradoxical function of TGF-β. To the best of our knowledge, this is the first report that shows significant gene - gene correlation in the TGF-β signaling pathway in patient-derived PDA based solely on tMUC1 expression levels. Most significant was the SMAD4 correlation to tMUC1, ID4, SRC, and CDKN2B in high versus low tMUC1 PDA. Another interesting pattern that emerged was the significant correlation of ID4 to tMUC1, and all other genes in tMUC1 high versus low PDA. These data indicate the clinical relevance of tMUC1 in modulating the TGF-β signaling in PDA. In addition to the bioinformatics data, we report significant correlation of tMUC1 to TGF-βRI, and RII protein levels in a panel of human PDA cell lines which informs the downstream signaling in response to TGF-β1. Thus, TGF-β1 induces phosphorylation of c-Src and activates the non-canonical Erk/MAPK pathway in the tMUC1-high PDA cells (which also expresses higher TGF-βRII). While, in the tMUC1-low cells (that expresses higher levels of TGF-βRI), TGF-β1 induces activation of the canonical pathway, SMAD2/3 phosphorylation, leading to increased apoptosis. Furthermore, we show that TGF-β1 regulates tMUC1 protein level in PDA cells by increasing proteasomal degradation of tMUC1 suggesting a possible feedback loop. Finally, *in vivo*, treatment with neutralizing antibody to TGF-β significantly reduced tumor progression and final tumor burden in MUC1-high PDAC tumors. In contrast the MUC1-low PDAC tumors did not respond at all to the treatment. Thus, we suggest that MUC1 expression may be used to personalize the treatment with TGF-β targeted treatment modalities.

## Acknowledgements

We thank Dr. Chandra Williams, DVM, DACLAM, CPIA, University Veterinarian and Vivarium Director and all vivarium staff members. Thanks to Benjamin Jaquez for help with Tissue Culture.

## REFERENCES

1. McGuigan, A., et al., Pancreatic cancer: A review of clinical diagnosis, epidemiology, treatment and outcomes. World journal of gastroenterology, 2018. 24(43): p. 4846–4861.

2. Ilic, M. and I. Ilic, Epidemiology of pancreatic cancer. World journal of gastroenterology, 2016. 22(44): p. 9694–9705.

3. Noone AM, H.N., Krapcho M, Miller D, Brest A, Yu M, Ruhl J, Tatalovich Z, Mariotto A, Lewis DR, Chen HS, Feuer EJ, Cronin KA (eds). SEER Cancer Statistics Review, 1975-2015, National Cancer Institute. 2018; based on November 2017 SEER data submission, posted to the SEER web site, April 2018.]. Available from: https://seer.cancer.gov/csr/1975_2015/.

4. Namwanje, M. and C.W. Brown, Activins and inhibins: roles in development, physiology, and disease. Cold Spring Harbor perspectives in biology, 2016. 8(7): p. a021881.

5. Isabel, F., et al., TGF-beta Signaling in Cancer Treatment. Current Pharmaceutical Design, 2014. 20(17): p. 2934–2947.

6. Carcamo, J., et al., Type I receptors specify growth-inhibitory and transcriptional responses to transforming growth factor beta and activin. Mol Cell Biol, 1994. 14(6): p. 3810–21.

7. Shi, Y., Structural insights on Smad function in TGFβ signaling. BioEssays, 2001. 23(3): p. 223–232.

8. Colak, S. and P. ten Dijke, Targeting TGF-β Signaling in Cancer. Trends in Cancer, 2017. 3(1): p. 56–71.

9. Massague, J., TGFbeta in Cancer. Cell, 2008. 134(2): p. 215–30.

10. Mittal, V., Epithelial Mesenchymal Transition in Tumor Metastasis. Annual Review of Pathology: Mechanisms of Disease, 2018. 13(1): p. 395–412.

11. Massague, J. and Y.G. Chen, Controlling TGF-beta signaling. Genes Dev, 2000. 14(6): p. 627–44.

12. Wrana, J.L., et al., Mechanism of activation of the TGF-β receptor. Nature, 1994. 370(6488): p. 341–347.

13. Tsukazaki, T., et al., SARA, a FYVE Domain Protein that Recruits Smad2 to the TGFβ Receptor. Cell, 1998. 95(6): p. 779–791.

14. Eppert, K., et al., MADR2 Maps to 18q21 and Encodes a TGFβ–Regulated MAD– Related Protein That Is Functionally Mutated in Colorectal Carcinoma. Cell, 1996. 86(4): p. 543–552.

15. Lee, M.K., et al., TGF-beta activates Erk MAP kinase signalling through direct phosphorylation of ShcA. Embo j, 2007. 26(17): p. 3957–67.

16. Schutte, M., et al., DPC4 gene in various tumor types. Cancer Res, 1996. 56(11): p. 2527–30.

17. Nath, S. and P. Mukherjee, MUC1: a multifaceted oncoprotein with a key role in cancer progression. Trends in Molecular Medicine. 20(6): p. 332–342.

18. Zhou, R., et al., A novel association of neuropilin-1 and MUC1 in pancreatic ductal adenocarcinoma: role in induction of VEGF signaling and angiogenesis. Oncogene, 2016.

19. Kufe, D.W., Mucins in cancer: function, prognosis and therapy. Nature reviews. Cancer, 2009. 9(12): p. 874–885.

20. Tinder, T.L., et al., MUC1 enhances tumor progression and contributes toward immunosuppression in a mouse model of spontaneous pancreatic adenocarcinoma. J Immunol, 2008. 181(5): p. 3116–25.

21. Roy, L.D., et al., MUC1 enhances invasiveness of pancreatic cancer cells by inducing epithelial to mesenchymal transition. Oncogene, 2011. 30(12): p. 1449–59.

22. Kato, K., et al., MUC1: The First Respiratory Mucin with an Anti-Inflammatory Function. J Clin Med, 2017. 6(12).

23. Singh, P.K and M.A. Hollingsworth, Cell surface-associated mucins in signal transduction. Trends in Cell Biology, 2006. 16(9): p. 467–476.

24. Thompson, E.J., et al., Tyrosines in the MUC1 cytoplasmic tail modulate oncogenic signaling pathways. Cancer Research, 2005. 65(9 Supplement): p. 222–222.

25. Galliher, A.J and W.P. Schiemann, Src Phosphorylates Tyr&lt;sup&gt;284&lt;/sup&gt; in TGF-β Type II Receptor and Regulates TGF-β Stimulation of p38 MAPK during Breast Cancer Cell Proliferation and Invasion. Cancer Research, 2007. 67(8): p. 3752.

26. Grover, P., et al., SMAD4-independent activation of TGF-beta signaling by MUC1 in a human pancreatic cancer cell line. Oncotarget, 2018. 9(6): p. 6897–6910.

27. Sahraei, M., et al., MUC1 regulates PDGFA expression during pancreatic cancer progression. Oncogene, 2012. 31(47): p. 4935–4945.

28. Besmer, D.M., et al., Pancreatic ductal adenocarcinoma mice lacking mucin 1 have a profound defect in tumor growth and metastasis. Cancer Res, 2011. 71(13): p. 4432–42.

29. Elston, R. and G.J. Inman, Crosstalk between p53 and TGF-β Signalling. Journal of signal transduction, 2012. 2012.

30. Adorno, M., et al., A Mutant-p53/Smad complex opposes p63 to empower TGFβ-induced metastasis. Cell, 2009. 137(1): p. 87–98.

31. Ihle, N.T., et al., Effect of KRAS oncogene substitutions on protein behavior: implications for signaling and clinical outcome. Journal of the National Cancer Institute, 2012. 104(3): p. 228–239.

32. Simeone, D.M., T. Pham, and C.D. Logsdon, Disruption of TGFβ signaling pathways in human pancreatic cancer cells. Annals of surgery, 2000. 232(1): p. 73.

33. Subramanian, G., et al., Targeting endogenous transforming growth factor β receptor signaling in SMAD4-deficient human pancreatic carcinoma cells inhibits their invasive phenotype 1. Cancer research, 2004. 64(15): p. 5200–5211.

34. Grover, P., et al., SMAD4-independent activation of TGF-β signaling by MUC1 in a human pancreatic cancer cell line. Oncotarget, 2018. 9(6): p. 6897.

35. Roskoski Jr, R., Src protein–tyrosine kinase structure and regulation. Biochemical and biophysical research communications, 2004. 324(4): p. 1155–1164.

36. Roskoski Jr, R., Src kinase regulation by phosphorylation and dephosphorylation. Biochemical and biophysical research communications, 2005. 331(1): p. 1–14.

37. Irtegun, S., et al., Tyrosine 416 is phosphorylated in the closed, repressed conformation of c-Src. PLoS One, 2013. 8(7).

38. Jang, H.H., Regulation of protein degradation by proteasomes in cancer. Journal of cancer prevention, 2018. 23(4): p. 153.

39. Lomako, W.M., et al., TGFbeta regulation of membrane mucin Muc4 via proteosome degradation. J Cell Biochem, 2009. 107(4): p. 797–802.

40. Cheever, M.A., et al., The prioritization of cancer antigens: a national cancer institute pilot project for the acceleration of translational research. Clinical cancer research, 2009. 15(17): p. 5323–5337.

41. Ozdamar, B., et al., Regulation of the polarity protein Par6 by TGFß receptors controls epithelial cell plasticity. Science, 2005. 307(5715): p. 1603–1609.

42. Valderrama-Carvajal, H., et al., Activin/TGF-β induce apoptosis through Smad-dependent expression of the lipid phosphatase SHIP. Nature cell biology, 2002. 4(12): p. 963–969.

43. Rape, M. and S. Jentsch, Taking a bite: proteasomal protein processing. Nat Cell Biol, 2002. 4(5): p. E113–6.

44. Ahmed, S., et al., The TGF-β/Smad4 signaling pathway in pancreatic carcinogenesis and its clinical significance. Journal of clinical medicine, 2017. 6(1): p. 5.

45. Derynck, R., R.J. Akhurst, and A. Balmain, TGF-β signaling in tumor suppression and cancer progression. Nature genetics, 2001. 29(2): p. 117–129.

46. Miyazono, K. and K. Miyazawa, Id: a target of BMP signaling. Sci. STKE, 2002. 2002(151): p. pe40–pe40.

47. Chan, A.S.W., et al., Downregulation of ID4 by promoter hypermethylation in gastric adenocarcinoma. Oncogene, 2003. 22(44): p. 6946–6953.

48. Umetani, N., et al., Epigenetic inactivation of ID4 in colorectal carcinomas correlates with poor differentiation and unfavorable prognosis. Clinical cancer research, 2004. 10(22): p. 7475–7483.

49. Wheeler, D.L., M. Iida, and E.F. Dunn, The role of Src in solid tumors. The oncologist, 2009. 14(7): p. 667.

50. Li, Y., et al., The c-Src tyrosine kinase regulates signaling of the human DF3/MUC1 carcinoma-associated antigen with GSK3β and β-catenin. Journal of Biological Chemistry, 2001. 276(9): p. 6061–6064.

51. Wen, Y., et al., Nuclear association of the cytoplasmic tail of MUC1 and β-catenin. Journal of Biological Chemistry, 2003. 278(39): p. 38029–38039.

52. Pai, P., et al., Mucins and Wnt/β-catenin signaling in gastrointestinal cancers: an unholy nexus. Carcinogenesis, 2016. 37(3): p. 223–232.

53. Zhang, S., et al., RhoA regulates G1-S progression of gastric cancer cells by modulation of multiple INK4 family tumor suppressors. Molecular Cancer Research, 2009. 7(4): p. 570–580.

54. Doublier, S., et al., RhoA silencing reverts the resistance to doxorubicin in human colon cancer cells. Molecular Cancer Research, 2008. 6(10): p. 1607–1620.

55. Fu, L. and N. Kettner, Progress in molecular biology and translational science. 2013, Elsevier.

56. Wiggs, J.L., Glaucoma genes and mechanisms, in Progress in molecular biology and translational science. 2015, Elsevier. p. 315–342.

57. Raphael, B.J., et al., Integrated genomic characterization of pancreatic ductal adenocarcinoma. Cancer cell, 2017. 32(2): p. 185–203. e13.

58. Schroeder, J.A., et al., Transgenic MUC1 interacts with EGFR and correlates with MAP kinase activation in the mouse mammary gland. Journal of Biological Chemistry, 2001.

59. Grossman, R.L., et al., Toward a Shared Vision for Cancer Genomic Data. New England Journal of Medicine, 2016. 375(12): p. 1109–1112.

60. McKenzie, A.T., et al., DGCA: A comprehensive R package for Differential Gene Correlation Analysis. BMC systems biology, 2016. 10(1): p. 106–106.

61. Murphy, A.M., et al., Vesicular stomatitis virus as an oncolytic agent against pancreatic ductal adenocarcinoma. J Virol, 2012. 86(6): p. 3073–87.

62. Massagué, J. and D. Wotton, Transcriptional control by the TGF‐β/Smad signaling system. The EMBO journal, 2000. 19(8): p. 1745–1754.

63. Ross, S. and C.S. Hill, How the Smads regulate transcription. The international journal of biochemistry & cell biology, 2008. 40(3): p. 383–408.

64. Bunda, S., et al., Src promotes GTPase activity of Ras via tyrosine 32 phosphorylation. Proceedings of the National Academy of Sciences, 2014. 111(36): p. E3785–E3794.

65. Fey, D., et al. The complexities and versatility of the RAS-to-ERK signalling system in normal and cancer cells. in Seminars in cell & developmental biology. 2016. Elsevier.

