## Supplemental Figure 1 for "Tumor -Associated MUC1 Regulates TGF-β Signaling and Function in Pancreatic Ductal Adenocarcinoma"

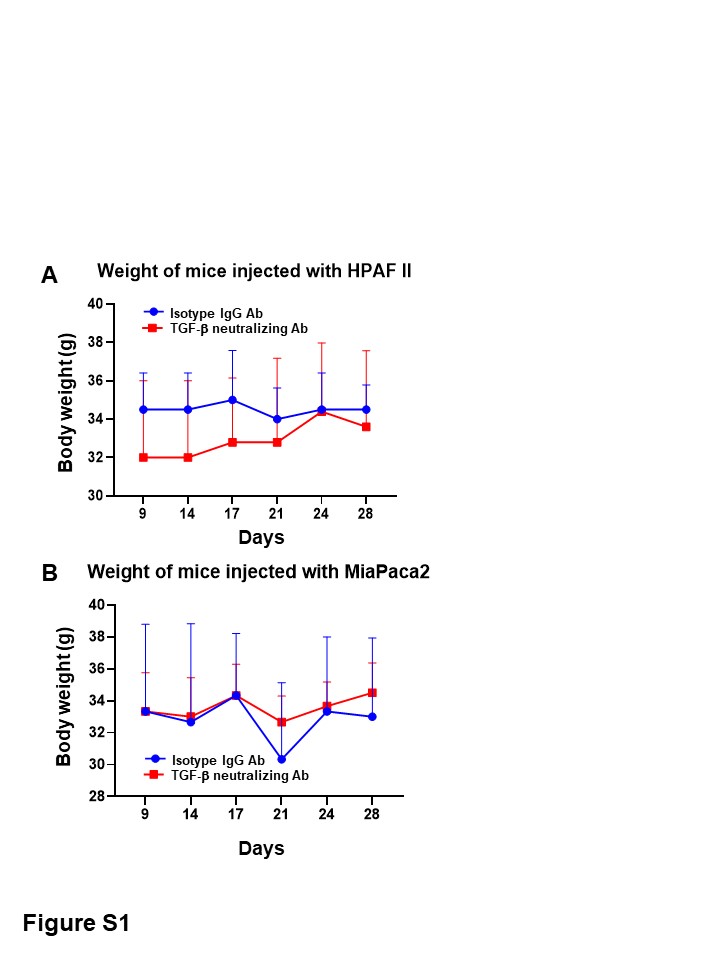

**Supplemental Figure 1: Neutralizing TGF-β1 antibody or IgG isotype antibody treatment did not have any adverse effect on the weight or well-being of the mice. A.** Weight of mice with HPAFII tumors (n=5 for TGF-β neutralizing Ab and n=4 for IgG isotype) and **B.** Weight of mice with MiaPaca2 (n=6 for both groups).
